# Precise replacement of *Saccharomyces cerevisiae* proteasome genes with human orthologs by an integrative targeting method

**DOI:** 10.1101/2020.03.25.006262

**Authors:** Christopher M. Yellman

## Abstract

Artificial induction of a chromosomal double-strand break in *Saccharomyces cerevisiae* enhances the frequency of integration of homologous DNA fragments into the broken region by up to several orders of magnitude. The process of homologous repair can be exploited to integrate, in principle, any foreign DNA into a target site, provided the introduced DNA is flanked at both the 5’ and 3’ ends by sequences homologous to the region surrounding the double-strand break. We have developed a tool set that requires a minimum of steps to induce double-strand breaks at chromosomal target sites with the meganuclease I-SceI and select integration events at those sites. We demonstrate this method in two different applications. First, the introduction of site-specific single-nucleotide phosphorylation site mutations into the *S. cerevisiae* gene *SPO12*. Second, the precise chromosomal replacement of eleven *S. cerevisiae* proteasome genes with their human orthologs. Placing the human genes under *S. cerevisiae* transcriptional control allowed us to update our of model of functional replacement. Our experience suggests that using native promoters may be a useful general strategy for the coordinated expression of foreign genes in *S. cerevisiae*. We provide an integrative targeting toolset that will facilitate a variety of precision genome engineering applications.

## INTRODUCTION

The integration of DNA into *Saccharomyces cerevisiae* chromosomes has become a foundational tool for the creation of inheritable modifications of many types, including gene-epitope fusions, mutations, and foreign gene insertions. DNA transformed into *S. cerevisiae* can integrate stably into chromosomes by homologous recombination when it has sequence homology to the target site^1–3^. Linear double-stranded DNA integrates more efficiently than circular DNA, and can carry heterologous DNA into the integration site as a consequence of recombination at the DNA ends.

The presence of a double-strand break (DSB) at the target site further increases the efficiency of DNA integration by homology-directed repair (HDR)^4^. The experimental induction of DSBs to initiate recombination at specific sites was pioneered in *Saccharomyces cerevisiae* using the HO meganuclease^5^, followed soon after by the I-*Sce*I meganuclease^6^. Meganucleases have since been used in *S. cerevisiae*, other microbes and even metazoan species to enhance the efficiency of chromosomal modifications^6–10^. In principle, a variety of meganucleases will work in yeast, but I-*Sce*I^11,12^ and I-*Cre*I^13^ have been the most frequently used. The “*delitto perfetto”* is a particularly elegant method that uses I-SceI for DSB induction and scarless repair with templates as small as oligonucleotides^12,14–16^. More recently the RNA-guided endonuclease Cas9 has become a widely-used tool for DSB induction in yeast^17–22^ and other organisms^23^.

The meganucleases have large DNA recognition sequences, usually 18-24 base pairs (bp) long, so they are unlikely to occur randomly in the relatively small genomes of yeast. The use of a meganuclease in genome engineering therefore requires that its recognition sequence be integrated at or near the target site to prepare it for DSB induction. In contrast, Cas9 can be directed to a large variety of target sites using unique guide RNAs (gRNAs). However, several considerations affect the utility of CRISPR-Cas9 for editing yeast genomes, and suggest that meganucleases will continue to be useful.

Firstly, it is difficult to predict the efficiency of DSB induction by Cas9 at specific gRNA sites. Factors that inhibit the performance of individual gRNAs include the presence of nucleosomes at the target site^24^ and intrinsic sequence features of the RNA^25^. As a result, several gRNA candidates must often be compared experimentally to find one that performs with high efficiency^26^. Secondly, good gRNA targets, while numerous in *S. cerevisiae*^17^, are not ubiquitous. Consequently, the use of oligonucleotides, which are potentially very useful repair templates, is limited to chromosomal sites with an efficient gRNA target in the region spanned by the oligonucleotide. Thirdly, a gRNA target that is not fully disabled by the DSB repair continues to be available for repeated cutting, potentially biasing the repair towards undesired events. Fourthly, CRISPR-Cas9 has well-documented off-target effects that continue to be actively investigated^27,28^, although they are of less concern in yeast than in organisms with larger genomes. Finally, when using CRISPR-Cas9, a specific repair event can be selected from all possible events only if it confers a novel selectable phenotype. When the DSB is induced within an essential gene, the selection for repair to a viable state is strong^29^, but changes made at non-essential loci lack that advantage.

We have developed a simplified method for genome engineering *S. cerevisiae* using I-SceI for DSB induction. While conserving the key features of “*delitto perfetto”*, we have reduced the cassettes for DSB induction and +/- selection from ∼ 4.6 kb to less than 1.3 kb, and provided a variety of separate plasmid-borne or integrated constructs for I-SceI expression. Our integrative targeting (*IT*) cassettes carry only a single marker, *K. lactis URA3*, and built-in I-SceI recognition sites at one or both ends. We used the *IT* method to introduce phosphorylation site mutations into the gene *SPO12* from oligonucleotide repair templates, and to precisely replace essential yeast proteasome genes with their human orthologs. Placing human proteasome orthologs under *S. cerevisiae* transcriptional control allowed us to refine our understanding of cross-species complementation by human proteasome subunits in yeast. Our methods outline a high-confidence work flow for genome engineering of *S. cerevisiae*, and we provide a variety of strains that are useful starting points for further applications.

## MATERIALS AND METHODS

### Plasmids

#### Plasmids carrying IT cassettes

The *IT* cassettes (Figure 1) were synthesized by PCR using plasmid pOM42^31^ as the template for the *Kluyveromyces lactis URA3* gene, including 299 bp of its native promoter and 117 bp of its terminator. I-SceI recognition sequences were incorporated, in various orientations, into the PCR primers used to amplify *K. lactis URA3*. The PCR products were cloned by Cold Fusion (SBI) into a plasmid backbone derived from pGEM-7Zf(+) to make the *IT* plasmids (Table 1).

**Table 1.**
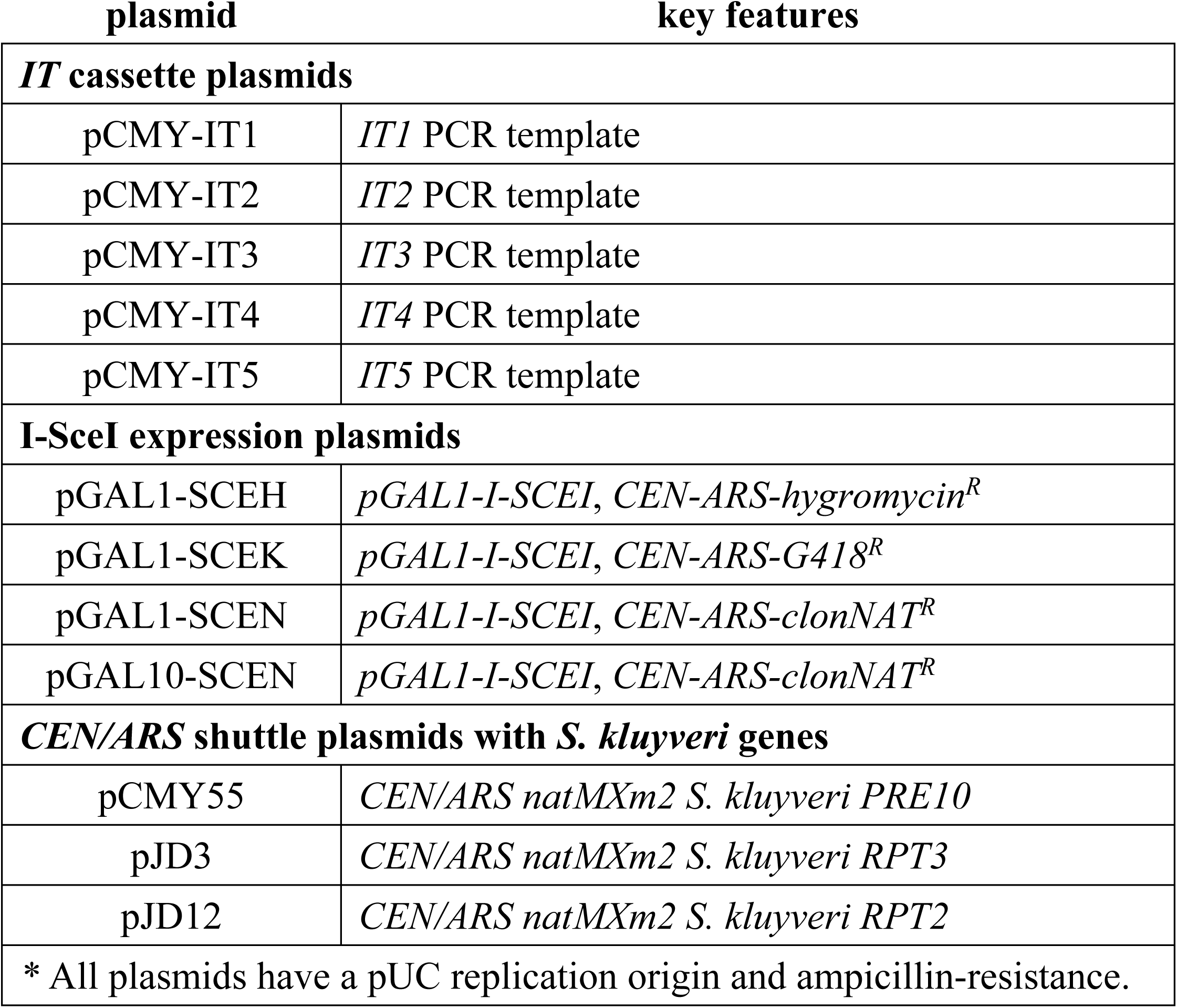
Plasmids made or used in this study*.

**Figure 1.**
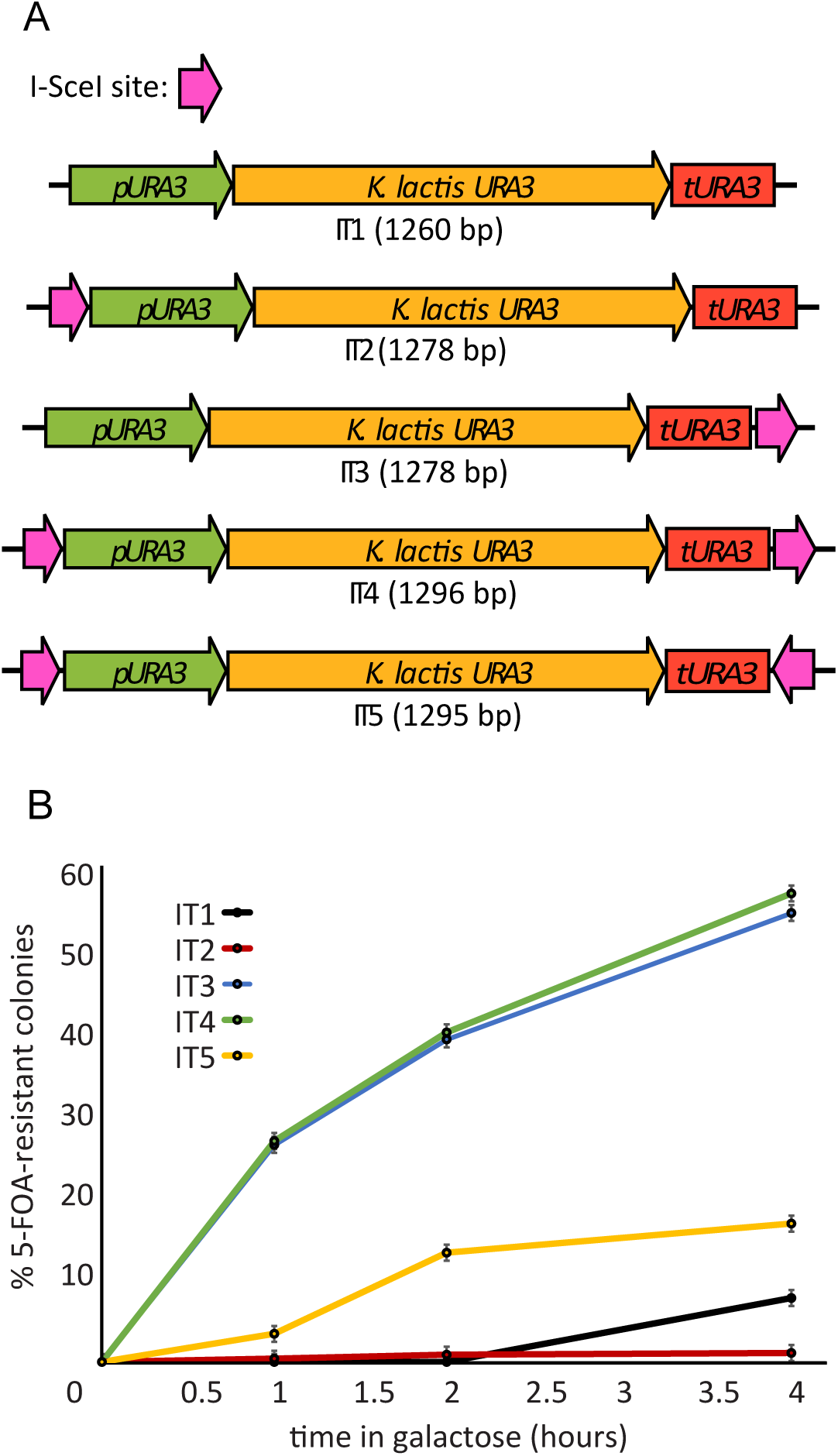
*IT* cassettes are targets for double-strand break induction by I-SceI. (A) Cassettes *IT1-IT5* contain the *K. lactis URA3* gene, integral I-SceI recognition sites in various orientations, and common PCR priming sites at their 5’ and 3’ ends. (B) Formation of 5-FOA-resistant colonies following induction of double-strand breaks at cassettes *IT1-IT5*. In diploid yeast, the *IT* cassettes were chromosomally-integrated into one copy of chromosome IV at the site *GT2* (Table 2, strains CMY 3427, 3428, 3429, 3021, 3430). The homologous chromosome was unmodified. A single copy of p*GAL1-I-SCEI* chromosomally-integrated at *GT1* was used to induce I-SceI expression upon the addition of galactose. Cells were sampled at the indicated time points by plating on YPD for single colonies, then replica-plating to SC -Ura + 5-FOA plates to count the fraction of 5-FOA-resistant cells in the population. At least 84 cells of each strain were analyzed at time zero, and the number cells counted increased to more than 300 of each strain by the 4-hour time point. Error bars represent the standard error of the mean calculated from three separate platings of cells from the same culture.

#### Plasmids for I-SceI expression

*pGAL1-I-SCEI* expression modules were assembled in the yeast *CEN/ARS* plasmid backbones pRS41H, pRS41K and pRS41N^32^ by *in vivo* homologous recombination in *S. cerevisiae*. The backbones were linearized with the endonuclease EcoRV, then co-transformed into yeast with three PCR products consisting of a 503 bp *GAL1* promoter from pYM-N22^33^, the *I-SCEI* open reading frame^12^, and a 201 bp *S. cerevisiae* native *GAL1* terminator. The PCR fragments had overlapping homology of ∼ 45 bp at each junction to drive their assembly. The assembled plasmids were recovered by preparation of yeast genomic DNA and electroporation into *E. coli*, and sequenced across the assembled regions. Plasmid pGAL10-SCEN was assembled in the pRCVS6N (CMY, unpublished) backbone with a 480 bp *S. cerevisiae* native *GAL10* promoter and a 136 bp *GAL10* terminator

**Table 2.**
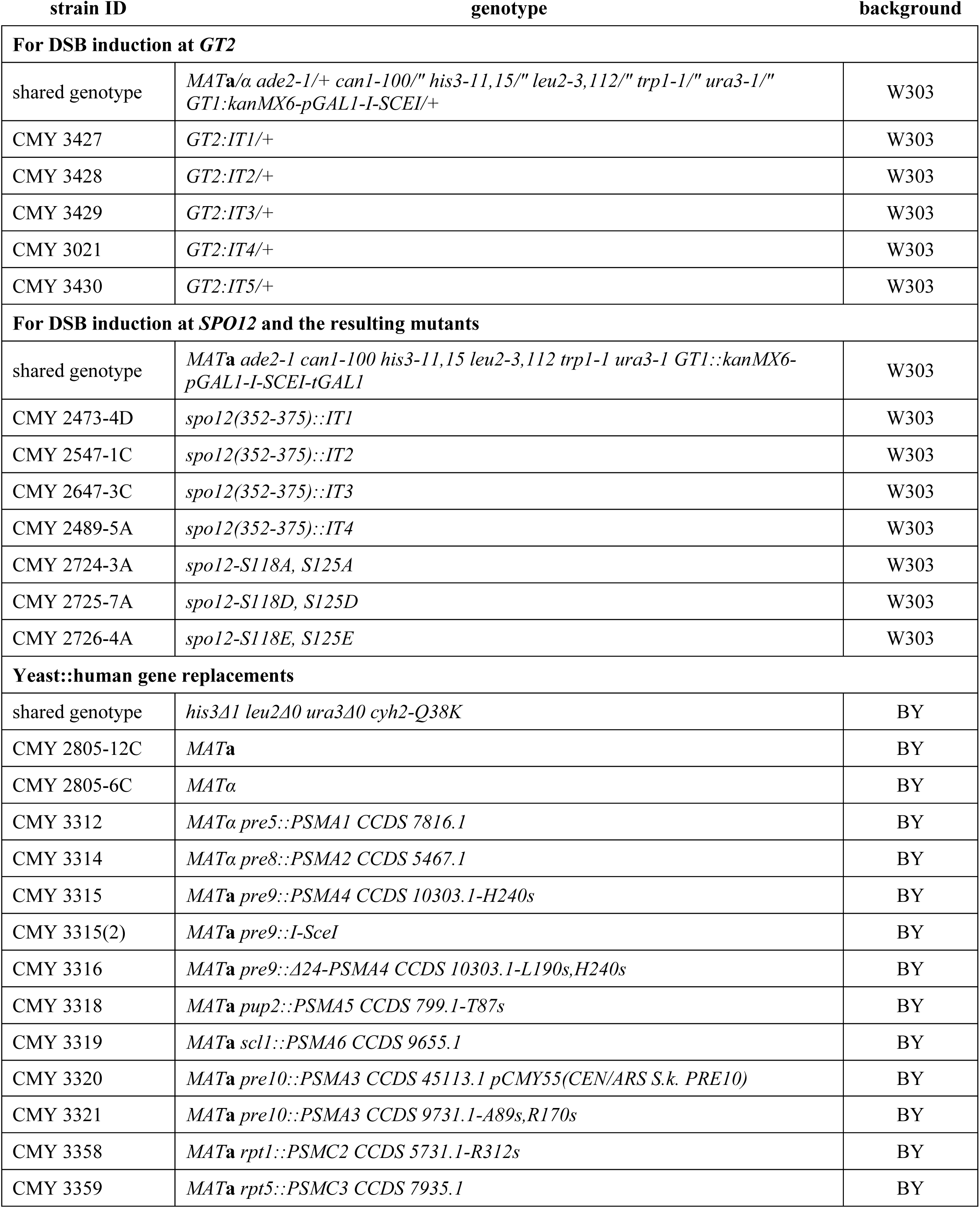
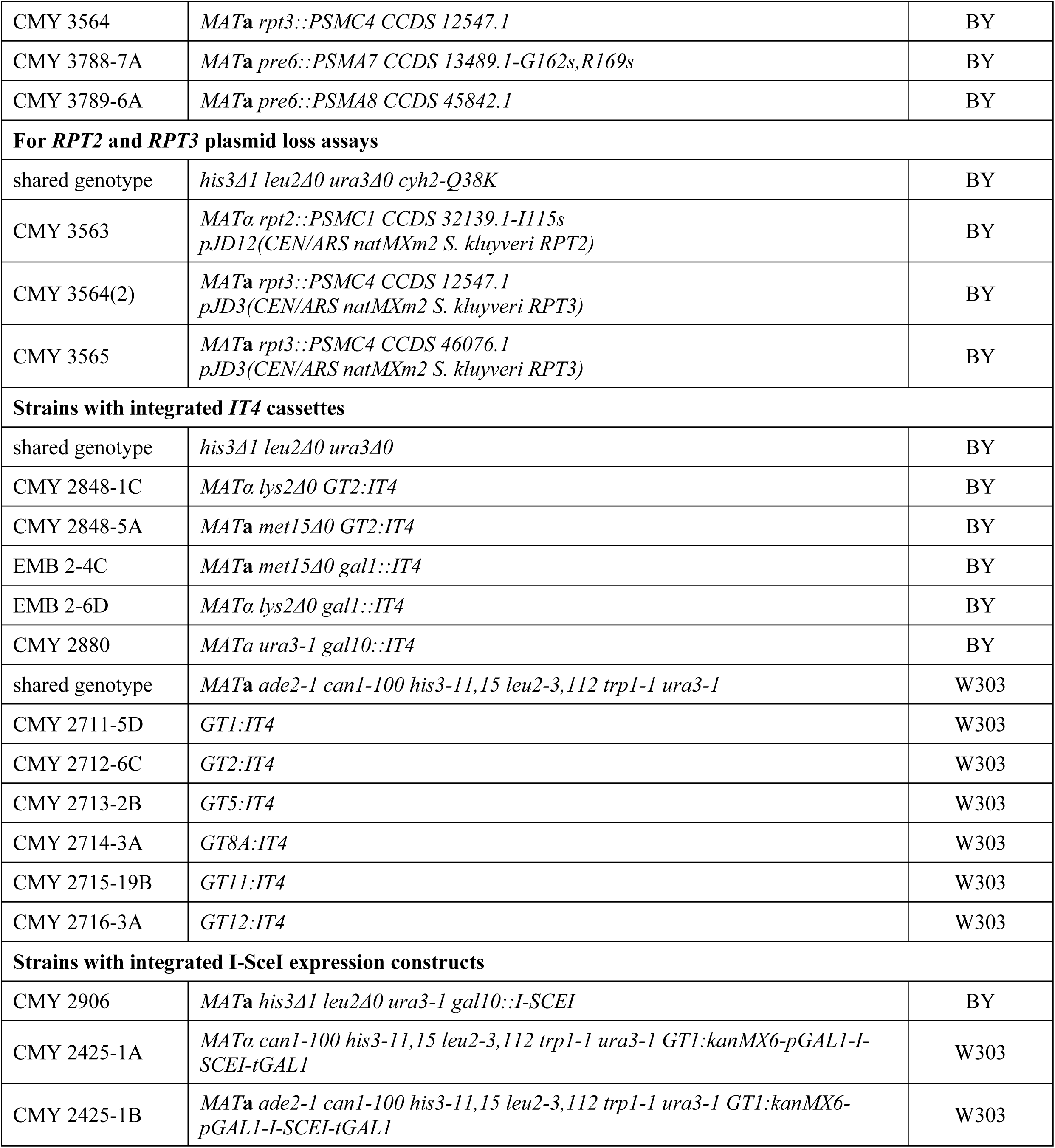
*S. cerevisiae* strains used in this study.

*Plasmids for complementation of S. cerevisiae gene deletions with S. kluyveri genes* Complementing plasmids carrying *S. kluyveri* orthologs of *S. cerevisiae* proteasome genes have been previously described^34^.

### Amplification and chromosomal integration of *IT* cassettes

Chromosomal integration of an *IT* cassette requires its synthesis with PCR primers that have priming regions common to all of the cassettes and unique identity to the 5’ and 3’ regions flanking the desired integration site. Integration of the cassettes into a chromosomal site is relatively efficient when the flanking target identity at each end is at least 40 bp. The forward primer (5’ to 3’) requires 5’ target identity + CGGACGTCACGACCTGCG and the reverse primer (5’ to 3’) requires 3’ reverse complementary identity + GGCTGTCAGGCGTGCACG. Recommended PCR amplification conditions are described in Table S1.

### Yeast media, DSB induction and transformation

Yeast media and growth conditions were standard^35^.

I-SceI expression was induced in yeast cells from the *GAL1* or *GAL10* promoters as follows: Cells were grown overnight in YP/2% raffinose, inoculated at ∼1 × 10^6^ cells/ml into fresh YP/2% raffinose in the morning, and grown for 3-4 hours to ensure they were in logarithmic growth. At the zero time point of I-SceI expression, galactose was added to the cycling cells to reach a final concentration of 2%. Induction continued for different lengths of time depending on the experiment.

DNA transformations into yeast were performed using the PEG/lithium acetate high-efficiency method^36^. The typical transformation targeted ∼1 × 10^8^ yeast cells.

### Yeast strains

Yeast strains were all of the BY or W303 backgrounds. Strain names and genotypes are listed in Table 2. All viable strains with chromosomal modifications were backcrossed to a congenic strain, and derivatives of either mating type are available upon request.

### Identification of neutral genomic target (*GT*) sites

By inspection of chromosomal sequences from the *Saccharomyces* genome database (SGD), we identified a set of genomic targets in *S. cerevisiae* to use for the integration of I-SceI expression constructs and as general sites for the integration of foreign DNA. Their chromosomal locations are summarized in Table S2. We did not work with all of the *GT* sites, but include their locations for potential use.

### Oligonucleotides

Double-stranded oligonucleotides were prepared by mixing single-stranded oligonucleotides together at a concentration of 50 µM each in 10mM Tris, pH 8.0/50 mM NaCl. The mixture was heated at 95°C in a heat block for 10 minutes, and cooled to room temperature over a period of approximately one hour to promote annealing.

### Human gene coding sequences

The coding sequences for human open reading frames (ORFs) were amplified by PCR from plasmids in the human ORFeome collection (hORFeome V7.1), with the exception of *PSMA8 CCDS 45842*.*1*, which was amplified from plasmid HsCD00336796 (Harvard Institute for Proteomics).

### DNA sequencing

All plasmids were confirmed by Sanger sequencing of at least the relevant assembled construct. All chromosomally-integrated constructs, including *IT* cassettes, I-SceI expression modules, *SPO12* mutations and human ORFs were sequenced after integration. The loci were amplified by PCR from outside the regions of yeast sequence identity used for homologous recombination, and sequenced across the entire construct.

## RESULTS

### Minimal integrative targeting (*IT*) cassettes with +/- selection

We constructed integrative targeting cassettes containing only the marker gene *Kluyveromyces lactis URA3*, which can be both positively and negatively selected, and recognition sequences for the homing endonuclease I-SceI (Figure 1). The set of cassettes includes versions that contain no I-*Sce*I site at all, a single site at either the 5’ or 3’ end of the cassette, or sites at both the 5’ and 3’ ends, in direct or inverse orientation to each other. The cassettes are maintained on high-copy *E. coli* plasmids (Table 1) that serve as PCR templates.

The cassettes, amplified by PCR with flanking target identity, can be integrated into a yeast chromosomal target locus by high-efficiency transformation^36^ and selected for by complementation of a *ura3* mutation in the host strain. The eventual replacement of the cassette with a DNA cargo is selected for using media containing 5-fluoro-orotic acid (5-FOA), which is lethal to Ura+ yeast^37^ that have not excised or mutated the *K. lactis URA3* gene.

### Control of I-*Sce*I expression

For flexible control of DSB induction, we generated constructs that place I-SceI expression under the control of the strongly repressible and inducible *S. cerevisiae GAL1* and *GAL10* promoters. Yeast centromeric plasmids carrying I-SceI expression constructs (Table 1) can be transformed into yeast and selected with a variety of dominant drug-resistance markers, then spontaneously lost during unselected growth. We also provide several yeast strains with *pGAL1* or *pGAL10*-driven I-SceI expression from chromosomally-integrated constructs (Table 2).

Expression of I-*Sce*I can therefore be controlled using a variety of methods appropriate to different applications.

### The positions and orientations of I-SceI sites affect the efficiency of HDR

We wanted to measure the efficiency with which the five *IT* cassettes, when integrated into a chromosomal site, would induce homologous recombination. To best estimate the frequency of chromosomal DSB formation at the different cassettes, we designed an assay in which the repair template was supplied on a homologous chromosome, and therefore available as efficiently as possible. In diploid cells, the *IT* cassettes were integrated into one copy of chromosome IV at a neutral genomic locus we refer to as *GT2* (Table 2). Following DSB induction, repair of the break using the homologous chromosome eliminated the *IT* cassette and the cells became Ura- and 5-FOA-resistant (5-FOA^R^). With the homologous chromosome being present in every cell, the rate of recovery of FOA^R^ cells should quantitatively reflect DSB formation.

We induced I-SceI expression in diploid cells and counted 5-FOA^R^ colonies as a fraction of the total at 0, 1, 2 and 4-hour intervals (Figure 1), expecting that the presence of two I-SceI sites instead of one would increase DSB formation. To our surprise, cassettes *IT3* and *IT4* performed equally well, yielding 5-FOA^R^ in close to 60% of all cells after 4 hours in galactose. In contrast, *IT1, IT2* and *IT5* induced 5-FOA^R^ poorly. Because the DSB induction results were unexpected, we confirmed the identities of the *IT* cassette strains by PCR length analysis of the 5’ and 3’ ends of the cassettes at all time points in the experiment (data not shown). The most parsimonious interpretation of the data is that the forward-oriented I-SceI site at the 3’ end of *IT3* and *IT4* performs far better than any other break site. The forward-facing I-SceI site at the beginning of *IT1* is a weak DSB site, while the inversely-oriented I-SceI sites in *IT5* together are relatively inefficient. Although we think it is unlikely, we cannot rule out the possibility that local features affected the DSB activity induced in the experiment, and therefore that the cassettes might perform differently in other chromosomal contexts.

### Chromosome engineering applications

To explore the utility of the *IT* cassettes for genome engineering, we performed two types of chromosomal modifications. The first was the introduction of phosphorylation site mutations into the non-essential *SPO12* gene using double-stranded oligonucleotides as the repair templates. The second set of modifications was the precise replacement of eleven yeast genes encoding subunits of the proteasome with the coding sequences of their human orthologs.

### Repair of *spo12::IT* with double-stranded oligonucleotides to create phosphorylation site mutations

Spo12 protein is an activator of the early anaphase release of the phosphatase Cdc14, and serines 118 and 125 of Spo12 are required for this release^38^. We used the *IT* method to make inhibitory and activating phosphomimetic mutations at these two amino acid residues. An S to A (alanine) mutation approximates a serine that cannot be phosphorylated, while mutations to D (aspartic acid) and E (glutamic acid) mimic phosphorylated serine^39^. To introduce the mutations, the *IT1, IT2, IT3* and *IT4* cassettes were first integrated into the non-essential *SPO12* gene, replacing twenty-four base pairs (bp 352-375) (Figure 2). We induced DSBs in the *IT* cassettes and transformed the cells with double-stranded oligonucleotides to repair the breaks.

**Figure 2.**
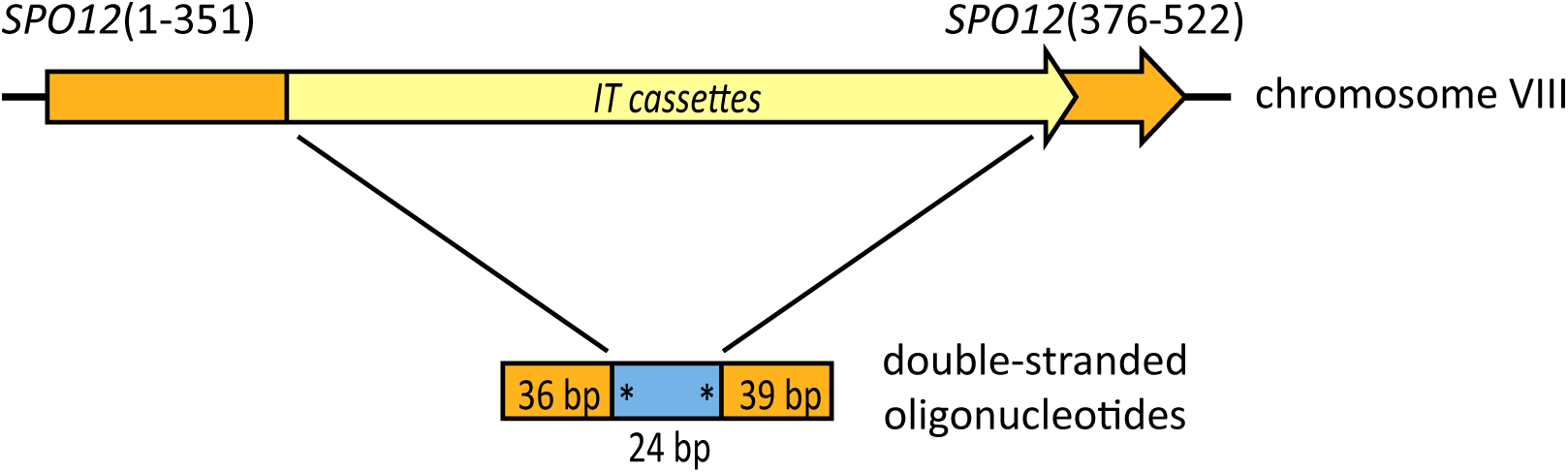
DSB repair with oligonucleotides creates phosphorylation site mutations in *SPO12*. In haploid yeast, cassettes *IT1-IT4* were integrated into the *SPO12* gene, replacing base pairs 352-375 (Table 2, strains CMY 2473-4D, 2547-1C, 2647-3C and 2489-5A). I-SceI expression was induced from chromosomally-integrated p*GAL1-I-SCEI* for 90 minutes, at which point the cells were transformed with double-stranded oligonucleotides. The mutagenized sites are indicated with asterisks.

The relative efficiencies of the four different *spo12::IT* cassettes as targets for repair with an oligonucleotide encoding S118A/S125A were consistent with the results of our assay of DSB repair at *GT2* (Table 3). The *spo12::IT1* target formed 5-FOA^R^ colonies inefficiently and was a poor repair site. *spo12::IT2* yielded relatively few 5-FOA^R^ colonies, but most of them used the oligonucleotides for repair. The *IT3* and *IT4* versions formed 5-FOA^R^ colonies efficiently and consistently used the oligonucleotides as repair templates. We used *spo12::IT4* to introduce three pairs of mutations (S118A/S125A, S118D/S125D and S118E/S125E), each time recovering ∼3000 colonies of which 30/30 were repaired as desired.

**Table 3.** *spo12(352-375)::IT* cassette replacement efficiencies.

| **target cassette** | **# of FOA<sup>R</sup> colonies** | **FOA<sup>R</sup> efficiency** | **correct repair rate** |
| --- | --- | --- | --- |
| *spo12:IT1* | 53 | $\sim 5.3 \times 10^{-7}$ | 2/10 |
| *spo12:IT2* | $\sim 250$ | $\sim 2.5 \times 10^{-6}$ | 9/10 |
| *spo12:IT3* | $\sim 2300$ | $\sim 2.3 \times 10^{-5}$ | 10/10 |
| *spo12:IT4* | $\sim 3000$ | $\sim 3 \times 10^{-5}$ | 30/30 |

In summary, repair of DSBs induced at *IT3* and *IT4* yielded many candidates and consistently used the desired oligonucleotide templates. DSBs induced at *IT3* were repaired using oligonucleotides that spanned ∼1.3 kb from the break site, consistent with the previously reported oligonucleotide-templated repair of DSBs induced at one end of the ∼4.6 kb CORE cassette^4^.

### Replacement of essential yeast proteasome genes with their human orthologs

The eukaryotic proteasome is a highly conserved protease with approximately 30 protein subunits, responsible for the degradation of ubiquitinated proteins^40,41^. We have previously shown, in plasmid-based complementation tests, that many human genes encoding subunits of the proteasome can functionally replace their yeast orthologs under the control of a strong constitutive yeast promoter and terminator^30^. However, such assays are affected by plasmid instability and copy number variation and the need to grow the cells in selective media. The ability of a heterologous gene to support viability is also subject to the activity level of the chosen promoter, a variable which is often not well understood.

#### A gene replacement strategy to protect genetic stability

The proteasome has direct roles in chromosome segregation^42^ and DNA double-strand break repair^43,44^. Therefore, we designed a work flow of several high-confidence steps that minimized the risk of genotoxic stress on the yeast cells due to partial or temporary loss of proteasome activity (Figure 3). We first transformed diploid yeast to replace one copy of each gene with the *IT4* cassette. Diploid yeast heterozygous for the *gene::IT4* deletions were then transformed with centromeric plasmids carrying the orthologous *Saccharomyces kluyveri* gene, under the control of the *S. kluyveri* promoter and terminator, which we have previously shown are able to complement the *S. cerevisiae* gene deletions^34^. The diploid cells were sporulated and tetrads dissected to recover haploid cells with *gene::IT4* deletions covered by the plasmid-borne *S. kluyveri* genes. The *IT4* cassettes were then replaced by inducing I-SceI and transforming with PCR-amplified human ORFs, flanked by homology to the promoter and terminator of the yeast gene. Because standard 60-mer PCR primers were used, the regions of flanking homology were relatively short, ranging from 32-44 NT at the 5’ ends and 27-41 NT at the 3’ ends, with one exception that had slightly longer homology.

**Figure 3.**
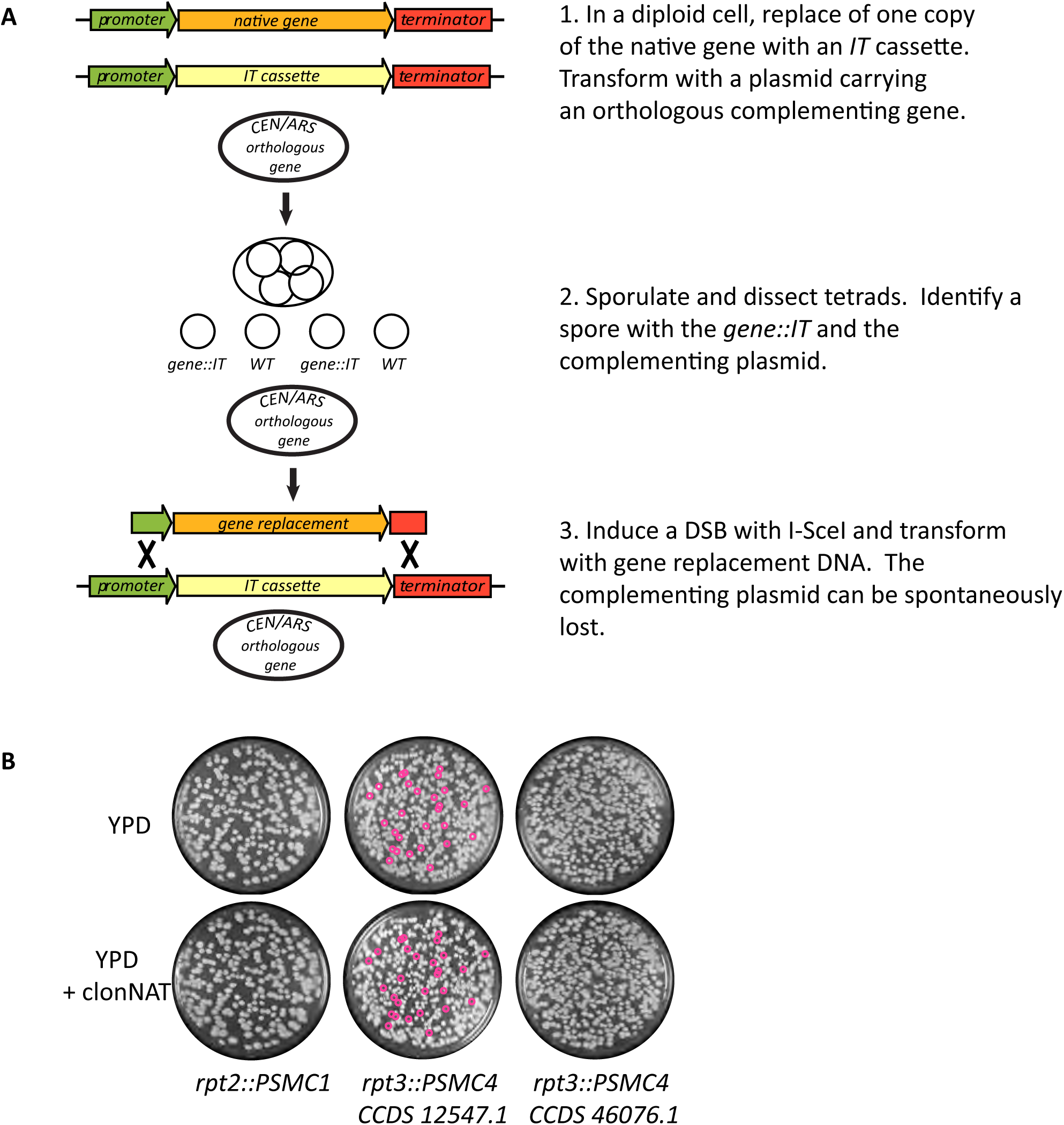
Replacing essential *S. cerevisiae* genes and testing complementation. (A) An essential gene is targeted for replacement in a diploid yeast cell, and function is maintained throughout the procedure by a complementing plasmid-borne orthologous gene. (B) Yeast (CMY 3563, 3564 and 3565) with human replacements of *RPT2* and *RPT3* were complemented by a plasmid carrying *S. kluyveri RPT2* (pJD12) or *RPT3* (pJD3). Cultures were grown in YPD liquid overnight to permit spontaneous plasmid loss, diluted and plated at low density to allow the formation of individual colonies. Plasmid loss was assayed by replica-plating to YPD + clonNAT. Small fuchsia circles indicate colonies that formed on YPD but not on YPD + clonNAT, indicating they had lost the complementing plasmid.

We isolated and screened 5-FOA^R^ gene replacement candidates by yeast colony PCR and sequenced a variety of them. In addition to recovering candidates with the desired repair to human ORFs, we found two types of undesired repair products, namely mutations in *K. lactis URA3* and deletions that reduced the *IT4* cassette to a single, unmarked I-SceI site. Analysis of 252 candidates showed that 38 (15%) had repaired to the human ORF, 31 (12%) had mutated *URA3* and 183 (73%) had reduced the cassette to an I-SceI site. The I-SceI site reductions must have occurred by microhomology-mediated end joining, a relatively high-fidelity form of repair in yeast^45,46^ We did not observe this type of repair product when replacing the *SPO12* targets with oligonucleotides. Their high frequency in the PCR product transformations underscores the importance of efficient delivery of the repair template. The PCR products we used to introduce human ORFs were both larger and less abundant than the oligonucleotides used to introduce mutations into *SPO12*. Therefore, the efficiency of transformation was probably much lower with the large DNA fragments. Cassette reduction will usually be an undesired outcome, but it can be avoided by using *IT3*, which carries only a single I-SceI site.

We confirmed the human ORFs integrations by sequencing the chromosomal loci from outside the regions of homology present in the PCR products and across each integrated human ORF. As we had hoped, we found no cases in which the orthologous *S. kluyveri* genes were used as templates for repair of the induced DSBs, due to the sequence divergence of their promoters and terminators.

#### Functional complementation by chromosomally-integrated human genes

To test the ability of human proteasome genes to function when controlled by the native yeast promoters and terminators, we precisely replaced eleven yeast proteasome genes, at their chromosomal loci, with the orthologous human protein coding sequences. The genes we replaced encoded the seven subunits of the core α ring and four of six subunits of the ATPase ring, which is responsible for substrate translocation into the catalytic core^41^. The resulting yeast strains each contained a single human coding sequence.

Many human proteasome genes have splicing isoforms, so we compared all human cDNA source clones to the consensus coding sequences (CCDS) in the NCBI Gene database, and identified the CCDS most similar to the clones. We used PCR to modify several of the cDNAs so that their sequences and lengths matched at least one CCDS in the database, leaving only a few instances of silent nucleotide changes. Table S3 summarizes the yeast:human orthology relationships of the proteasome core and ATPase ring subunits we worked with, the length and sequence comparisons of our cDNA clones to the database CCDS, and the existence of splicing isoforms.

Among the human CCDS used for gene replacements, there was at least one isoform of each core α-ring subunit and three out of four ATPase subunits that minimally supported viability on rich medium (Figure 4). In a previous study, we reported that the human α2 (Psma2) subunit was toxic when expressed from a strong constitutive yeast promoter.^34^. Placing it under fully native yeast transcriptional control allowed it to support viability. We did not previously test complementation by the α4 human paralogs Psma7 and Psma8, due to the lack of cDNA clones, but we now done so. Psma7 is a widely-expressed isoform, while Psma8 is testis-specific and has three isoforms of different lengths. Both Psma7 and the mid-length isoform of Psma8 supported viability (Table S3). We did not test the short and long isoforms of Psma8. Not all of the human α-ring isoforms that we tested were able to complement yeast, however. Of the two splicing isoforms of Psma3, the human α7, only the longer one supported viability.

**Figure 4.**
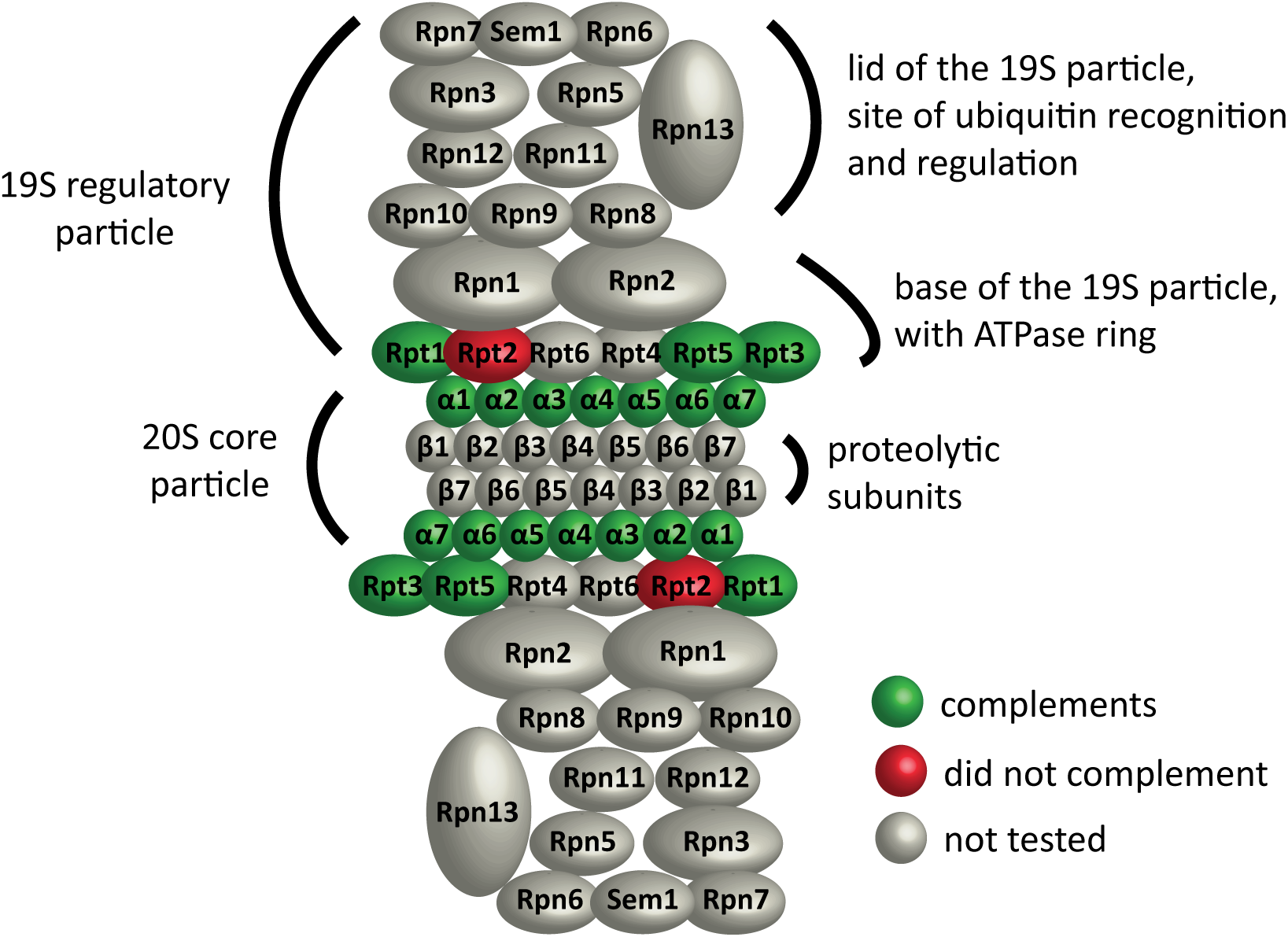
Functional replacement of yeast proteasome subunits with their human orthologs. Selected subunits of the proteolytic core α-ring and the ATPase ring of the yeast proteasome were replaced in their chromosomal loci with the orthologous human coding sequences. Human subunits that supported viability are green, those that did not are red and untested subunits are gray.

The ATPase ring consists of six subunits, all of which supported viability in our previous study^34^. We replaced subunits Rpt1, Rpt2, Rpt3 and Rpt5 with their human orthologs Psmc2, Psmc1, Psmc4 and Psmc3 respectively. Psmc2, Psmc4 and Psmc3 supported viability, but Psmc1 did not (Figure 4 and Table S3). Of the two splicing isoforms of Psmc4, only the longer one supported viability (Table S3). We tested *rpt2::PSMC1* and *rpt3::PSMC4* activity by the ability of the strains to lose a complementing plasmid carrying *S. kluyveri RPT2* or *RPT3* (Figure 4C), leaving the human ORF as the only source of protein. Neither Psmc1 nor the short isoform of Psmc4 conferred on cells the ability to lose the complementing plasmid. However, the long isoform of Psmc4 allowed plasmid loss, indicating functional replacement. We previously reported that Psmc1 could functionally replace Rpt2, but by this more rigorous assay, we found it did not.

#### Yeast with individual human proteasome genes grow well in mitosis, but are sensitive to proteotoxic stress

The lineages leading to *S. cerevisiae* and humans diverged approximately 1 billion years ago^47^, so it would be surprising if human proteasome subunits had fully normal activity in yeast. The proteasome is required for both routine mitotic division^48^ and degradation of misfolded proteins^49,50^, so we compared the growth of unmodified and humanized yeast under both permissive conditions and proteotoxic stress.

Under permissive growth conditions (rich medium, 30°C), only a few humanized strains grew slowly (Figure 5A). Human Psma7 and Psma8 significantly delayed growth, while Psma1 caused a slight delay. The yeast α3 subunit Pre9 is the only non-essential protein in the core α- ring. Deletion of Pre9 causes a slight growth delay^51^, an effect which was very subtle in our assay. Yeast with the human α3 subunit Psma4 grew normally, but the Δ24-Psma4 N-terminal truncation allele caused a slight growth delay. Finally, the viable human Rpt ATPase strains grew at strikingly normal rates. Overall, humanized strains executed mitosis well under these permissive conditions.

**Figure 5.**
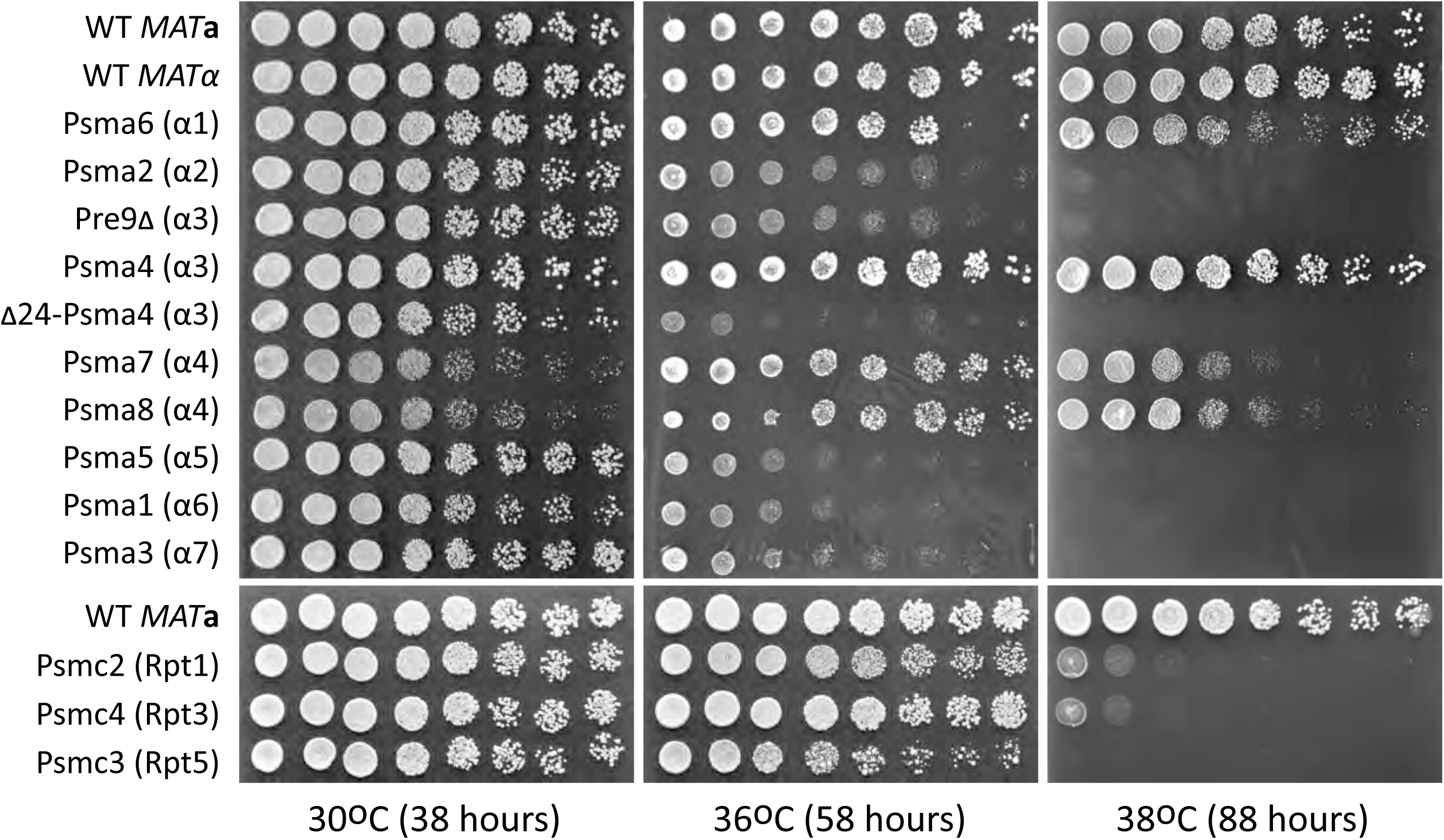
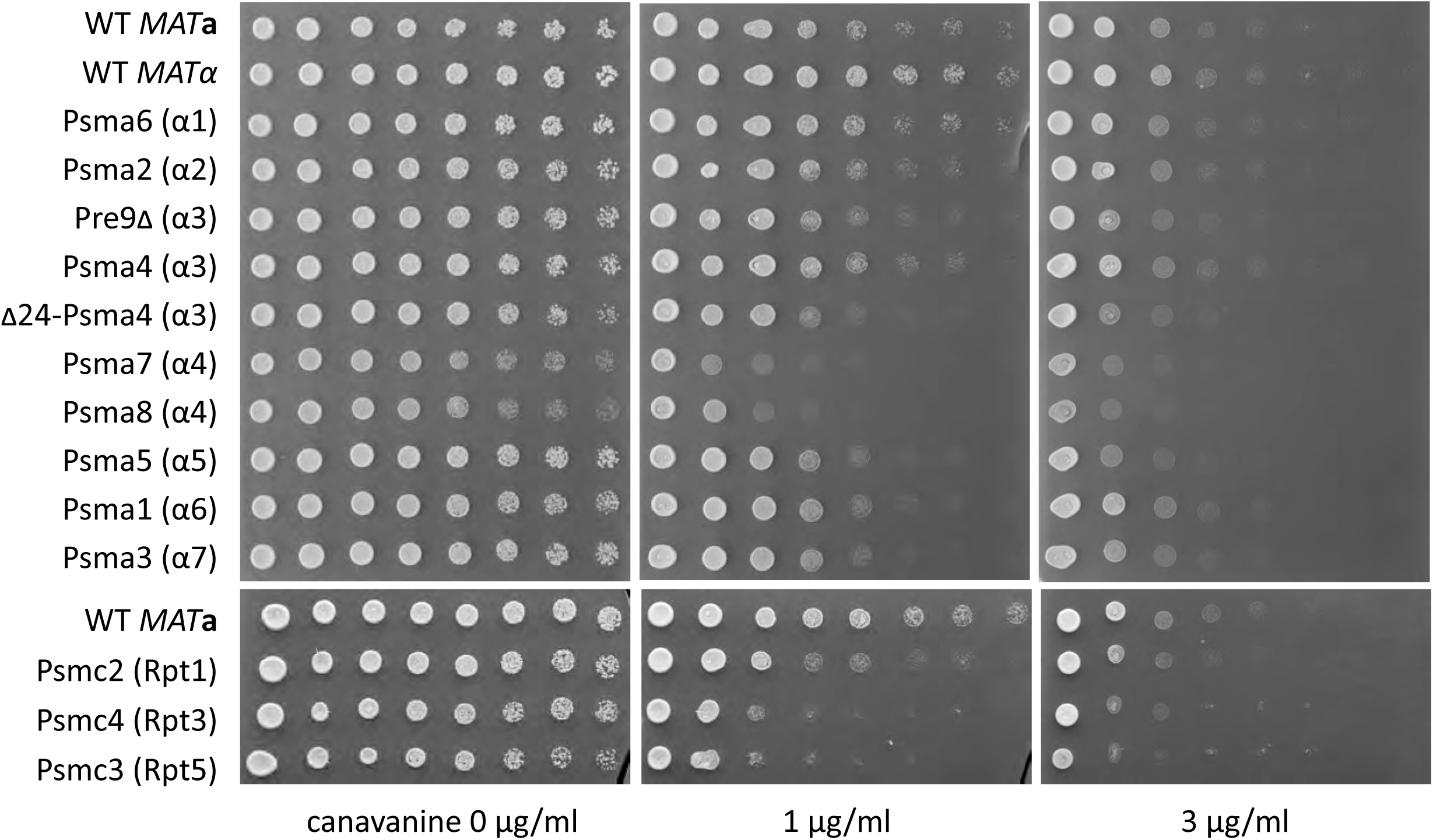
Yeast with individual human proteasome subunits are sensitive to proteotoxic stress. Yeast with individual human proteasome subunits (Table 2) were grown to saturation in liquid YPD medium at 30°C, then plated in 5-fold dilution series to assay growth. (A) Cells were plated on YPD and grown at 30°C for 38 hours, 36°C for 58 hours or 38°C for 88 hours. (B) Cells were plated on SC – Arg medium containing canavanine at 0, 1 or 3μg/ml and grown at 30°C for 40 hours.

High temperature imposes proteotoxic stress on cells in the form of misfolded proteins and survival of this stress requires the ubiquitin-proteasome system^52–56^. As previously reported, *pre9Δ* yeast were very sensitive to high temperature^57^ (Figure 5A). With a few exceptions, yeast with human proteasome subunits were also very sensitive to high temperature, growing poorly at 36°C and not at all at 38°C. The most striking exceptions were yeast with Psma6 (α1), which was slightly slow at 38°C and Psma4 (α3), which grew normally at 36°C and 38°C. The Psma7 (α4) and Psma8 (α4) strains grew fairly well at 36°C, and were only partially inhibited at 38°C. We conclude that most human substitutions compromise proteasome activity, but Psma1 and Psma4 provide almost normal activity at the elevated temperatures we tested.

Canavanine, a non-biogenic arginine analog, causes proteotoxic stress upon incorporation into new proteins, activating the yeast environmental stress response^58^. We performed a canavanine-sensitivity test on synthetic medium at 30°C to avoid temperature-dependent effects. In the absence of canavanine, the Δ24-Psma4 (α3), Psma7 (α4) and Psma8 (α4) strains grew slowly, as they had on YPD (Figure 5B). Yeast lacking Pre9 had been extremely sensitive to high temperature, but were only slightly sensitive to canavanine, suggesting the stress imposed by high temperature is more severe or general. The humanized α5, α6 and α7 strains (Psma5, Psma1 and Psma3) were slightly sensitive to canavanine. The humanized Rpt strains Psmc2, Psmc4 and Psmc3 were more strongly sensitive to canavanine, especially Psmc4 and Psmc3. In summary, our phenotypic analysis shows that yeast proteasomes with single human subunits tend to be severely deficient in the response to high temperature and moderately sensitive to protein misfolding.

## DISCUSSION

### Native transcriptional control is a powerful tool

Transcriptional circuitry is often complex, delicately balanced, and incompletely characterized. There is currently abundant interest in transferring complete foreign protein complexes and enzymatic pathways into yeast, for both functional conservation studies and synthetic biology applications. Native gene regulation may be particularly useful when working with complexes such as the proteasome, ribosome and CCT chaperonin, which have defined stoichiometries and are transcriptionally co-regulated^59–62^. Apart from these well-known examples, complexes formed during facultative responses, such as autophagy and DNA repair, are also co-regulated^61,63^, and the use of native promoters preserves those potentially important regulatory circuits.

Chromosomal integration of human proteasome genes under native transcriptional control allowed us to refine the complementation status of several proteasome subunits previously reported in a large-scale study^34^. In exceptional cases, altered expression may be desirable, but as a general strategy, native expression appears to be the best way to minimize confounding effects.

We benefitted from native gene regulation in one other way. From previous work, we knew that *S. kluyveri* proteasome genes, under native promoters and terminators, tend to complement *S. cerevisiae* gene deletions^34^. Having used these genes as part of our chromosomal integration strategy, we can now say, at least in a limited context, that the sequences of their promoters and terminators differ sufficiently from *S. cerevisiae* to make them unavailable for homologous repair. This trick won’t always work, but when available, it is very useful.

### We can learn from humanized yeast

We expected that human proteasome subunits, having evolved separately from yeast for 1 billion years^47^, would have pleiotropic functional deficits compared to native yeast subunits. The human subunits may be inefficiently synthesized or folded, they may limit the assembly or stability of the mature proteasome, or they may have more specific functional defects. Humanized yeast grew surprisingly well under permissive conditions, suggesting the proteasome was working well within its capacity, but phenotypic assays confirmed that the human genes did not provide fully normal activity. High temperature and protein folding stress exposed the incompleteness of the complementation. By exploring phenotypes ranging from minimal complementation to stress resistance, we were able to characterize the levels of functional conservation of human splicing isoforms and paralogs. In well-characterized cases of complementation, humanized yeast may be a useful platform to test the functional activity of human mutants, isoforms or splicing variants, or the efficacy of a drug treatment.

High temperature has wide-ranging effects on cells, including changes in membrane composition^64,65^, and the induction of multiple transcriptional and metabolic pathways^66,67^. Ubiquitin ligases and deubiquitinases of the ubiquitin-proteasome system are required for the degradation of proteins that misfold as a result of high temperature^55,56^. With the exception of the non-essential α3 subunit Pre9^57^, the roles of the proteasome core subunits in surviving high temperature have not been deeply investigated, in part because they are essential. The human substitutions can be can be viewed as temperature-sensitive alleles of essential proteasome subunits, but as with any classic temperature-sensitive or pleiotropic allele, there are a variety of possible reasons for the loss of function. Irrespective of these considerations, the failure to grow indicates that the core proteasome is required for survival at high temperature. The human substitutions may be useful reagents with which to further investigate the roles of the proteasome in high temperature growth and under proteotoxic stress.

### IT vs CRISPR/Cas9: pros and cons

Where are the pros and cons of the *IT* method and CRISPR/Cas9 in yeast genome engineering? Aside from the differences in DSB induction, CRISPR/Cas9 and *IT* have similar requirements in the subsequent steps of chromosomal modification. A repair template must be available during or soon after the DSB is made, and the desired repair products must be identified from a variety of possible events. Therefore, the main differences are in the convenience and level of control of DSB induction.

CRISPR/Cas9 is a very flexible way to induce chromosomal DSBs; sgRNA targets are abundant, and no preliminary chromosomal modification is necessary. This is particularly advantageous when working with essential genes. However, the performance of a specific CRISPR/Cas9 sgRNA is still unpredictable, and DSB induction near a specific locus typically requires comparing several sgRNA candidates. Therefore, CRISPR/Cas9 is ideal for large, exploratory experiments in which the efficiency of DSB induction at any one sgRNA target does not need to be optimal^26^. Some of the drawbacks of CRISPR/Cas9 can be overcome by offering large DNA repair templates, which span a chromosomal region containing multiple sgRNA sites.

The *IT* cassettes, on the other hand, must be integrated into a target site, and if that locus is essential for viability, the function must be covered, at least temporarily, with a complementing construct. *IT* cassettes can, in theory, be integrated into any chromosomal locus, and once integrated, DSB induction is, in our experience, independent of the locus. The freedom to choose a precise DSB induction site is potentially powerful, enabling intense mutagenesis of short, defined genomic regions using relatively cheap oligonucleotide repair templates.

One potential application of the *IT* method that we have only briefly explored is the integration of a cassette into the *kanMX4* marker used in the yeast standard gene deletion collection^68,69^. The systematic integration of an *IT* cassette into *kanMX4* would convert the gene deletion collection into a DSB induction collection. This and other applications will continue to make I-SceI a useful tool for yeast genome engineering.

## Supporting information

Supplementary Manuscript

## ACKNOWLEDGEMENTS

Thanks to Andreas Matouschek for his support during the preparation of this manuscript and to Joseph DeSautelle, who helped with plasmid construction.

Development of the IT method was started while CMY was a post-doctoral fellow of the Howard Hughes Medical Institute.

