## Supplementary material for "Precise replacement of *Saccharomyces cerevisiae* proteasome genes with human orthologs by an integrative targeting method": Figure S1

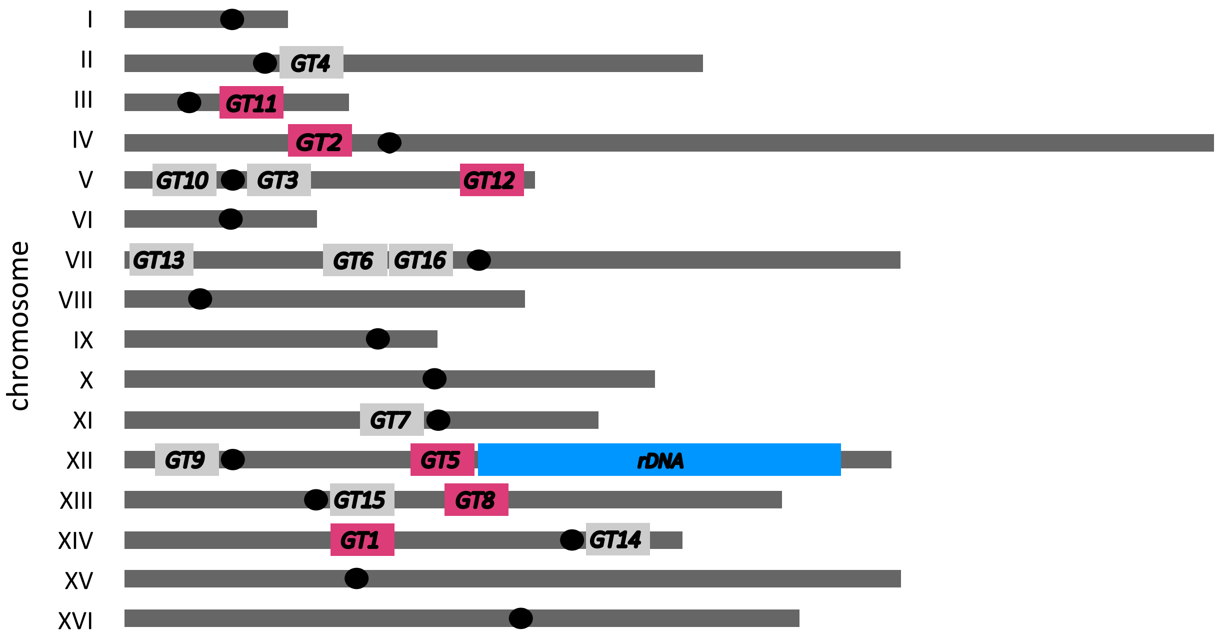


**Figure S1. Genomic targets (GTs): chromosomal loci amenable to the integration of foreign DNA.** Approximate locations of *GT* sites in the genome of *S. cerevisiae*. Chromosomes lengths are proportional to their actual sizes, and centromeres are marked with black ovals. The *IT4* cassette was integrated at six *GTs* (fuchsia) on different yeast chromosomes.
