## Supplementary Manuscript for "Precise replacement of *Saccharomyces cerevisiae* proteasome genes with human orthologs by an integrative targeting method"

**Yellman et al., supplemental materials**

**Additional *IT* resources**

During this project, we accumulated a variety of modified yeast strains that can be a “starter kit” for the *IT* method. We made yeast strains with *IT* cassettes integrated into a variety of chromosomal loci, including the native *GAL1* and *GAL10* genes, and identified a set of genomic targets we called *GTs* (Figure S1 and Table S2).

The chromosomal *gal1::IT* and *gal10::IT* constructs can be used as integration targets that place genes under *GAL* control, and their presence can be followed by a Gal^-^ phenotype. The stable expression of foreign genes under *GAL* control may be a useful for large-scale assays of conditional gene expression or overexpression phenotypes. When introducing foreign DNA into the *gal1::IT* and *gal10::IT* cassettes, it is important to express I-SceI from a construct that does not carry a repair template for the DSB site. For example, when replacing a *gal1::IT* cassette, I-SceI should not be expressed from a *pGAL1* construct, but from *pGAL10*. Conversely, when replacing *gal10::IT* cassettes, I-SceI should be under *pGAL1* control. The I-SceI expression plasmids (Table 1) provide flexibility to avoid this problem.

The sixteen potential genomic targets (*GTs*) that we identified should be amenable to the integration of foreign DNA, and we integrated *IT4* into six of them to prepare them for modification (Figure S1 and Table S2). In choosing sites, we sought to minimize the potential for interference with neighboring genes, promoters or other functional elements such as origins of replication. However, the ability of *S. cerevisiae* chromosomal sites to support heterologous expression can depend on local effects which are difficult to predict and best evaluated by the integration of a transcriptional reporter^1^. Our ability to successfully integrate *IT4* into all six *GT* sites we worked with indicates that, at those sites, the *K. lactis URA3* gene is at least sufficiently expressed to be selectable. The *GTs* are distributed throughout the *S. cerevisiae* genome, some with linkage to centromeres, telomeres or the rDNA array, but all from regions with actively transcribed nearby genes.
