## Supplementary material for "Precise replacement of *Saccharomyces cerevisiae* proteasome genes with human orthologs by an integrative targeting method": Table S1

**Table S1. Reaction conditions for PCR synthesis of *IT* cassettes.**

**Notes:**

The first few amplification cycles of the PCR synthesis use an annealing temperature appropriate to the common regions of all synthesis primers. Subsequent amplification cycles use a higher annealing temperature based on the T_m_ of the full-length primers. We show 70^o^C as the higher T_m_, but that value should be adjusted depending on the specific primer sequences used. The PCR products should be purified over a DNA minicolumn before being used in a high-efficiency PEG/LiOAc transformation of yeast.

predicted product sizes (base pairs):

*IT1*:	1260 + ~ 80 = ~ 1340

*IT2*, *IT3*:	1278 + ~ 80 = ~ 1358

*IT4*, *IT5*:	1296 + ~ 80 = ~ 1376

| **REACTION MIXTURE** | | | |
| --- | --- | --- | --- |
|  | **reagent** | **vol per**  **rxn (μl)** | **final**  **concentration** |
| **H_2_O** | pure water | 7.5 | - |
| **RXN mix** | 2x KAPA HiFi reaction mixture | 10 | 1x |
| **magnesium** | in reaction mixture | - | ? |
| **dNTPs** | in reaction mixture | - | ? |
| **polymerase** | in reaction mixture | - | - |
| **primers** | mixed primers, 3μM each | 2 | 0.3µM each |
| **template** | 20x-diluted plasmid DNA | 0.5 | - |
| **other reagent** |  |  |  |
| **each rxn** | complete reaction mixture | 20 | - |

| **THERMOCYCLER PROGRAM** | | | | |
| --- | --- | --- | --- | --- |
|  | **# of cycles** | **denaturation** | **annealing** | **synthesis** |
| **initial denaturation** | 1 | 96^o^C / 60s |  |  |
| **1^st^ amplification set** | 3 | 96^o^C / 20s | 56^o^C / 30s | 72^o^C / 75s |
| **2^nd^ amplification set** | 30 | 96^o^C / 20s | 70^o^C / 30s | 72^o^C / 75s |
| **final amplification** | 1 |  |  | 72^o^C / 120s |
| **store** | 1 |  | 4^o^C / hold |  |
