## Supplementary material for "Precise replacement of *Saccharomyces cerevisiae* proteasome genes with human orthologs by an integrative targeting method": Table S2

| **Table S2. Genomic target (*GT*) sites.** | | | |
| --- | --- | --- | --- |
| **locus** | **chromosome** | **SGD coordinates** | **comments** |
| *GT1* | XIV, left arm | 335522/335523 | *CBK1*-*YGP1* intergenic |
| *GT2* | IV, left arm | 443742/443743 | 5967 bp left of *CEN4; ATP16*-*MCD1* intergenic |
| *GT3* | V, right arm | 152859/152860 | 755 bp right of *CEN5* |
| *GT4* | II, right arm | 45436/45437 | 455 bp right of *CEN2* |
| *GT5* | XII, right arm | 449338/449339 | ~2.5kb left of the rDNA array |
| *GT6* | VII, left arm | 324086/324087 | *SRM1*-*TOS8* intergenic, tightly linked to *GT16* |
| *GT7* | XI, left arm | 439817/439818 | 311 bp left of *CEN11* |
| *GT8* | XIII, right arm | 512029/512030 | *NCW1-PKR1* intergenic |
| *GT9* | XII, left arm | 150359/150360 | 468 bp left of *CEN12* |
| *GT10* | V, left arm | 151409/151410 | 577 bp left of *CEN5* |
| *GT11* | III, right arm | 116855/116856 | 856 bp right of *CEN3, YCR001W-CDC10* intergenic |
| *GT12* | V, right arm | 555845/555846 | subtelomeric, *FAU1-TOG1* intergenic |
| *GT13* | VII, left arm | 16719/16720 | subtelomeric, *ADH4-ZRT1* intergenic |
| *GT14* | XIV, right arm | 629371/629372 | 496 bp right of *CEN14* |
| *GT15* | XIII, right arm | 268744/268745 | 595 bp right of *CEN13* |
| *GT16* | VII, left arm | 328174/328175 | *TOS8-VPS45* intergenic, in transposon *YGLWdelta6* |
